# Genomics and biogeography of novel *Trichodesmium* isolates from the Sargasso Sea

**DOI:** 10.64898/2026.07.30.741789

**Authors:** Catie S. Cleveland, Anjali M. Bhatnagar, Shelby J. Barnes, Yiming Zhao, Eric A. Webb

## Abstract

The cyanobacterium *Trichodesmium* is important in ocean nitrogen (N) and carbon (C) biogeochemistry through fixation of N_2_ gas and CO_2_ into NH_3_, amino acids, and carbohydrates. These essential compounds then fuel primary productivity, bacterial and grazing communities, and upper trophic levels. In the oligotrophic surface oceans, they are one of few taxa capable of utilizing both atmospheric C and N. *Trichodesmium* fall into 2 diazotrophic phylogenomic clades named “Thieb” and “Tery” after the most well characterized botanical species, *Trichodesmium thiebautii* and *Trichodesmium erythraeum*. Thieb is commonly the most abundant clade in all major ocean regions and paradoxically, is also the least characterized in terms of morphology, genomics, and ecophysiology due to underrepresentation in culture collections. In this study, 25 novel strains of Thieb were enriched in culture from the Sargasso Sea, phenotypically characterized, and whole genome sequenced. Cultures were found to belong to four well-supported subclades (i.e., ThiebA, ThiebB, ThiebC, and ThiebD). *Trichodesmium* subclades were found to have environmentally relevant differences in colony shape, gene content, cell dimensions, and inferred temperature range. Specifically, ThiebD and ThiebC had the largest biovolumes whereas ThiebA and ThiebB had the smallest. Importantly ThiebB also became relatively more abundant only in bulk seawater metagenomes collected at high temperature (>28°C). This finding indicates that as ocean warming continues, there may be shifts in *Trichodesmium* community structure towards smaller cells, which would affect overall C+N standing stocks.

## Introduction

*Trichodesmium* are globally important, biogeochemically relevant cyanobacteria which impact both the marine nitrogen (N) and carbon (C) cycle via nitrogen fixation and photosynthesis [1, 2]. *Trichodesmium* have large, cylindrical cells (∼5-30 µm diameter) which can connect into long (>100 µm) filaments that grow either as “free” planktonic forms or in colonies [3, 4]. Colonies generally fall into several different shapes, including “puffs” or “tufts,” which have filaments arranged radially or in parallel, respectively [1, 4]. N_2_-fixation carried out by *Trichodesmium* is an oxygen-sensitive process [5]. However, contrary to other filamentous diazotrophic cyanobacteria such as *Anabaena*, *Trichodesmium* have the unique ability to fix N_2_ gas and CO_2_ “simultaneously” using nitrogenase and RuBisCO without the use of differentiated heterocyst cells to protect the nitrogenase enzyme from O_2_ exposure [6, 7]. *Trichodesmium* grows in all major subtropical, oligotrophic ocean gyres, including the Indian Ocean, Atlantic Ocean/Sargasso Sea, and the Pacific Ocean [8–10]. In these locations, they can even form miles-long surface blooms, historically called “sea sawdust,” [1, 11–13] attesting to their large C + N standing stocks.

*Trichodesmium* are historically classified into 6 botanical species and two major clades using 16s-23s ITS and *hetR* sequence phylogenies, cell sizes, and pigmentation [3, 4, 14]. These species include *Trichodesmium erythraeum* and *Trichodesmium contortum* in clade I, and *Trichodesmium spiralis*, *Trichodesmium thiebautii*, *Trichodesmium tenue*, and *Trichodesmium hildebrandtii* in clade III and two other clades that have never been represented in culture collections (i.e., II and IV), [4]. Importantly, the *hetR* phylogenies of field and culture collections are what defined *Trichodesmium* into clade I, II, III, and IV [4]. More recent work has shown that the *hetR* gene can be paralogous, providing differing phylogenies in some strains, and is thus not an ideal phylogenetic marker in *Trichodesmium* [10, 15]. Based on recent whole genome sequencing efforts of cultures combined with increasing numbers of *Trichodesmium* environmental metagenome-assembled genomes (MAGs), it was determined that 16s-23s ITS phylogenetic clustering of clade I and clade III also corresponded to two major phylogenomic subclades [4, 10, 16, 17]. However, within each of the two phylogenomic clades, individual taxa have whole genome average nucleotide identity (ANI) >97% to each other, which is above the proposed inter-species cutoff of 95% [18]. This created difficulty in reconciling phenotypic botanical species designations with genomic sequences, particularly with environmental MAGs [10, 17]. Because of this phylogenetic uncertainty, isolate genomes and MAGs that cluster with clade I and clade III [4] were henceforth named as “Tery” and “Thieb,” respectively, after the most well-known botanical species of each clade (i.e., *T. erythraeum* and *T. thiebautii*).

Overall, Tery are generally found at much lower abundance in the oligotrophic ocean [8, 9, 16, 17], and based on their typically red, phycoerythrobilin-rich pigmentation, likely grow better in greenish, turbid waters rather than pure blue [4, 19]. Contrastingly, based on environmental qPCR surveys and metagenomic read recruitment, Thieb are significantly more numerically abundant (often ∼90% of the total *Trichodesmium* biomass in oligotrophic subtropical gyres), and have yellow-green, phycourobilin rich pigments better suited to clear, blue water [4, 8, 9, 16, 17]. Therefore, since Thieb dominate the oligotrophic oceans, which make up >60% of the marine environment [20], they are highly relevant to biogeochemical cycling and C+ N export.

Despite Thieb being most abundant, previous laboratory-based physiological studies have focused almost entirely on Tery (>300 studies of Tery vs approximately <5 for clade Thieb) due to underrepresentation of Thieb in culture. There was also previously only 1 whole genome sequence from Thieb, strain H9-4 [17], which is now extinct in culture. However, of the few strains that have been previously cultivated, Thieb members appear to exhibit colony type (puff vs tuft), cell diameter size, and 16S-23S ITS phylogeny diversity [4].

Herein, we introduce novel, diverse enrichments of Thieb isolated in artificial YBC II seawater media from the Sargasso Sea. Of these cultures, 25 were whole genome sequenced and 4 subclades, designated ThiebA, ThiebB, ThiebC, and ThiebD, were identified. These subclades had conserved differences in specialized metabolite, lipids, defense mechanism, and adhesion-related genes. Subclades also showed morphological diversity at the colony to the cellular level wherein ThiebD formed puffs or tufts and ThiebA only formed puffs with dense exopolysaccharides at the colony center. ThiebD also had the largest intracellular biovolume and fastest nitrogen fixation. The isolation of these strains therefore demonstrates that Thieb comprises many diverse, biogeochemically relevant members with unique subclade-specific features. Also, since Thieb are the most numerically abundant *Trichodesmium* group in the global oceans, characterization of their ecophysiology, ecology, morphology, and biogeographical range better constrains the impact *Trichodesmium* has on biogeochemical cycling in a changing global climate.

## Materials and Methods

### Isolation and Cultivation

Cultures were enriched from the euphotic zone near or at the Bermuda Atlantic Time Series site (BATS, 31.833°N, -64.167°W) in the Sargasso Sea and in the Gulf Stream (general location: 38.23, -69.91) aboard the R/V Atlantic Explorer over two years from June 2022-September 2022 and June 2023-September 2023. Colonies were obtained using a hand-towed 130-µm mesh plankton net (SEA-GEAR, Melbourne, FL, USA), hand-picked from the collected seawater using Pasteur pipetting, and rinsed twice in 0.2 µm-filtered natural seawater to remove larger eukaryotes and other loosely attached bacteria. Single, picked colonies were checked for any remaining attached eukaryotes on a Zeiss Stemi-4 dissecting microscope (Oberkoken, Germany) before being placed in ∼5 mL YBC II media [21] in sterile 14 ml polystyrene culture tubes (VWR^®^, Radnor, PA, USA). Cultures were incubated aboard the ship at ∼115 µmol Q m^-2^s^-1^ light level and 26°C in either flowing deck-board or portable electric incubators. The enrichments were subsequently transported back to the laboratory in insulated bags or boxes, and maintained in a Percival incubator (Perry, IA, USA) under the following conditions: YBC II media [21] amended with no vitamins, 12:12 diel cycle, ∼100-125 µmol Q m^-2^s^-1^ and 3000 K warm white lights, and 26°C. For the first month of initial cultivation, each culture was checked every 1-2 days and transferred to new media as needed. Enrichment cultures were acclimated to artificial YBC II seawater media and incubator conditions for ∼1 year. Select enrichment cultures (USCT24, USCT12, USCT02, and USCT15) were then further purified by diluting dense cultures to 1 filament per well in 96-well plates and re-sequencing the resulting biomass after cultures increased in density again. This was similar to the process by which strains VI-1, H-94, and others have been further purified following enrichment when filaments were separated by pipetting and re-grown from single filaments [4].

### DNA Extraction

*Trichodesmium* were grown in acid washed polycarbonate culture flasks or sterile polystyrene culture tubes prior to being gently vacuum filtered onto 5-µm polycarbonate filters, frozen, and used for DNA extractions. Between 10-100 mL were filtered for DNA extractions, depending on biomass and number of colonies. Genomic DNA was extracted using a modified phenol-chloroform-CTAB method [22, 23] amended with an overnight proteinase K (20 mg/mL stock; final 1.2 % : Sigma-Aldrich, Burlington, MA, USA), incubation at 55°C with SDS (25% stock; final 0.76 mg/ml), and final ethanol precipitation (if needed).

### Thieb Identification

*Trichodesmium rnpB* genes were amplified using both primer sets ten-*rnpB* and tery-*rnpB* designed specifically for Thieb and Tery, respectively [24]. Both primer sets have been found to have successful amplification of these two different clades [8, 9, 24]. The *rnpB* product sizes for Thieb vs Tery (102 vs 199 bp length) were visualized using ethidium bromide gel electrophoresis in 2% agarose and used to assign cultures to a broad clade prior to whole genome sequencing. Template DNA was also amplified from known strains of Thieb and Tery (*T. thiebautii* strain VI-I and *T. erythraeum* strain LIN [4] as positive controls for product size comparison.

### Genome Assembly and Taxonomy

Following extraction and identification using *rnpB* amplification, DNA was quantified and quality-checked using a Nanodrop 1000 spectrophotometer and whole genome sequenced at Novogene (Sacramento, CA, USA) using the Novaseq 6000 platform (Illumina, San Diego, CA, USA), 150 PE reads to generate raw forward and reverse reads (∼1 Gbp). These reads were trimmed using Trimmomatic v0.39 [25], quality verified using fastQC v0.11.9 [26], and assembled using metaSPAdes v4.2.0 [27, 28]. Binning was performed on contigs using Maxbin v2.2.7 [29], GTDB-tk v2.4.0 [30] was used to classify the bins, CheckM v1 and CheckM v2 [31, 32] were used to screen *Trichodesmium* bins for completeness and contamination, and FastANI v1.34 [18] was used to compute the average nucleotide identities of genomes. *Trichodesmium* bins were then imported into Anvi’o v8 [33] where they were further screen for contamination with Kaiju v1.10.1 [34], and any split sequences not belonging to phylum Cyanobacteria were then removed from each *Trichodesmium* genome. These genomes were then used to generate an initial *Trichodesmium* phylogenomic tree in GToTree [35] alongside *Trichodesmium* MAGs sequenced directly from oceanic samples [10, 17], (GCA 028982585.1, GCA 022448615.1, GCA 040262915.1, GCA 028982625.1, GCA 028982325.1, GCA 028982365.1) and cultivated Tery strains IMS101, LIN, and 21-75, and Thieb strain H9-4 (2023 version SAMN57478722). The initial tree was produced with 251 Cyanobacterial marker genes and filtered by only those having >50% of these genes per genome. The tree was rooted to *Okeania hirsuta* (GCA 003838195). The output was then passed to IQTree2 [36] and a final consensus tree was generated using 1,000 bootstraps and the best fit ModelFinder [37].

### Morphology and Microscopy

*Trichodesmium* cells were photographed at 100x on a Zeiss AxioStar microscope (Zeiss, Oberkochen, Germany) with phase contrast to measure and compare individual cell size, diameter, and appearance of cultures. All cultures were recently transferred within 7 days of imaging to ensure they were actively growing when imaged. Cell dimensions were measured using ImageJ v1 [38], and biovolumes were calculated assuming cylindrical-shaped cells as per previous methods [39]. Dark field images of whole colonies were also taken on a Zeiss STEMI-4 dissecting microscope (Zeiss, Oberkochen, Germany).

Scanning electron microscopy was performed on colonies that were first fixed in glutaraldehyde (∼2.5% in YBC II media), stored at 4°C, placed in microporous specimen capsules (Electron Microscopy Sciences, Hatfield, PA, USA), washed 1x in 0.1 M HEPES/5% sucrose/0.1% Triton-X buffer solution and 2x in ultrapure water with alternating microwaving at 250 watts for 1 minute each, and serially dehydrated from 50, 70, 95, 100% ethanol (EtOH) with microwaving at 250 watts for 1 minute between EtOH washes. The 100% EtOH washes were done 3x, and the last EtOH wash was performed with molecular sieves. The samples were then placed in a critical point drier (Tousimis, Rockville, MD, USA), mounted on SEM sample stubs (Ted Pella Inc., Redding, CA, USA) with carbon tape, and briefly coated with carbon. Upon first imaging, it was found that samples needed to be made more conductive, so they were additionally sputter coated with platinum and palladium. Images of whole colonies were taken on the Apreo 2S SEM (Thermo-Fisher, Waltham, MA, USA) with an Everhardt-Thornley detector, 1-2 kV accelerating voltage, and currents in the range of 25 pA-0.2 nA. Images of individual *Trichodesmium* and heterotroph cells were taken with the in-column T2 detector with OptiPlan mode and A+B, accelerating voltages ranging from 1-3 kV, and currents ranging from 6.3-25 pA.

### Pangenomic and Gene Functional Analyses

A pangenome was generated of the entire Thieb clade using the Anvi’o v8 pangenomic workflow (https://merenlab.org/2016/11/08/pangenomics-v2/) with an MCL threshold of 10 for highly similar genomes with high sequence similarity [33, 40]. Genomes were annotated using NCBI-COGs [41], Pfams with release 37.4 [42], KEGG-KOfam with release 2023-09-22, and HMMER v3.4 [43] to identify gene content differences across Thieb subclades. A FastANI heatmap was also created using the pangenome in Anvi’o to distinguish subclades by average nucleotide identity (ANI), and strains were ordered in the FastANI heatmap by gene cluster frequency, which groups strains by presence/absence of gene clusters and considers paralogous gene counts. The genomic “core” of the pangenome was defined as gene clusters shared between all strains in the pangenome, and subclade-specific accessory gene clusters are those shared by all strains within each individual subclade. Whole Thieb genomes were also annotated by AntiSMASH v8.0.4 [44] and BigSCAPE v2 [45] to identify specialized metabolite genes across subclades. AntiSMASH was used with Pfam v35.0 [42] and prodigal v2.6.3 [46] to first annotate biosynthetic gene clusters (BGCs), and BigSCAPE 2.0 [45] was used for clustering BGCs by similarity and presence vs absence across strains.

### *Trichodesmium* Subclade Biogeography

Habitat range was determined using metagenomic read recruitment to both *Trichodesmium* picked colonies [17] and GoShip bulk water reads collected using 0.22 µm filtration [47]. *Trichodesmium* picked colony metagenomes were previously sequenced along an East to West transect in the equatorial Atlantic Ocean [17] using the same methods as described above for culture colony collection with the following modifications: after dilution in 0.2 µm filtered seawater, ∼50 colonies were separated by “puff” and “tuft” morphologies, filtered down onto 5-µm polycarbonate filters and stored in liquid nitrogen until DNA extraction.

Whole genomes from individual Thieb subclades could not be used for read recruitment as the inter-subclade ANI values of the core genes were too similar. Therefore, using whole genomes as read recruitment targets would likely result in some amount of non-specific read mapping [48], which could skew abundance results. Therefore, the subclade-specific accessory genes were used as read recruitment targets instead. The following steps were taken prior to read recruitment: Thieb subclade specific accessory genes were identified using the pangenome workflow described previously, representative strain subclade specific accessory genes were exported with annotations, known mobile genetic element (MGE) or transposase-related genes were removed, BLAST+ v2.14.1 with BLASTX [49], (e-value cutoff of 1e-4) was used to identify and remove additional MGE genes not identified within Anvi’o [33, 50], and finally, the ultimate list of genes with a predicted function or conserved hypotheticals by BLASTX were retained and used for read recruitment. This allowed the accessory genes from representative strains of each subclade to serve as an abundance proxy for each unique taxonomic group.

Read recruitment to the picked colony and bulk water metagenomes was then performed using the following pipeline: metagenomic short reads were mapped using Bowtie v2.5.4 [51] to the conserved accessory genes of representative strains from each Thieb subclade as defined by the Anvi’o pangenomic workflow [33], Samtools v1.22.1 [52] converted Sequence Alignment Maps (SAMs) to Binary Alignment Maps (BAMs), CoverM v0.6.1[53] filtered BAMs at 99% identity over 85% of the read, Samtools v1.22.1 indexed the BAMs, Anvi’o profiled the reads, and finally, Anvi’o was used to calculate the mean coverage and detection values of each strain and each individual gene across all metagenomes. Mean coverage values across the accessory genes for representative strains of each subclade were converted to a “mean coverage ratio” to show relative abundance, and detection showed the percent of accessory genes with predicted functions for each strain/subclade that were at least 1x coverage.

For read recruitment results from GoShip 0.2 µm bulk water metagenomes, stations were plotted in a relative abundance bar plot with ggplot2 [54] where at least 1 Thieb subclade was found at >1x mean coverage. Thieb abundance data from these same stations were also used to calculate Bray-Curtis distances and to plot resulting values in an NMDS analysis in RStudio [55] using the “vegan” package [56]. K-means clustering was performed on the NMDS results, with an elbow plot being used to determine the number of k-means clusters used. A UMAP using the package “uwot” [57] and k-means clustering was plotted with the same stations, but abundance values were instead Z-scored prior to analysis. The NMDS and UMAP results were also plotted with station temperature ranges as well as by k-means clustering.

### Physiology

Nitrogen fixation was measured over the 12 daylight hours according to the acetylene reduction assay [58] with the following specifics: cultures were transferred 1:1 and incubated for 5 days, 30 mL aliquots of each strain were taken from polycarbonate culture flasks and placed in 43 mL Nalgene polycarbonate tubes with sealed septa caps, 2 mL of acetylene gas was added to the headspace of each tube, tubes were incubated at 26°C and ∼100 µmol Q/m^2^/s for 12 hours, and finally, ethylene production was measured following the daytime on a Shimadzu GC8-A (Shimadzu, Nakagyo-ku, Kyoto, Japan).

## Results and Discussion

### Cultures corresponded to the oligotrophic clade “Thieb”

Overall, >50 *Trichodesmium* enrichment cultures were collected from the Sargasso Sea and the Gulf Stream over summer-fall of 2022 and 2023. To start cultures, single colonies of *Trichodesmium* were placed individually in seawater media and incubated at 26 °C. Twenty-four of these new enrichment cultures were whole genome sequenced alongside one historical Thieb strain, VI-1 (**Supplemental Table S1**), which had known 16S-23S ITS and *hetR* gene sequences [4] but no full genome. Overall, 25 *Trichodesmium* strains and 217 associated bacterial MAGs were assembled in total and identified using GTDB-tk (**Supplemental Table S2**). Of these genomes, 49 of the *Trichodesmium*-associated heterotrophic bacteria corresponded to GTDB-tk RED novel genomes without contamination; this included 43 genomes with unknown species and 6 novel genera (**Supplemental Table S3**). The 25 *Trichodesmium* genomes were then compared to the previously sequenced genome of Thieb H9-4 (which is now extinct in culture). Together, these genomes were all conserved in GC content of ∼35% and were high in completion at ∼92-100% (**Supplemental Table S4**). The N50 values ranged from 10,028 to 22,891, and they had an average of ∼876 contigs across all genomes. These isolate genomes were then phylogenomically compared alongside high quality environmental MAGs to place them in an environmental context and determine subclade distinctions. From this it was found that the same overall clades “Thieb” and “Tery,” identified by 16S-23S ITS and 16S phylogenetics [4] were also present at the whole genome level. Within Thieb, the previously sequenced strain H9-4 grouped with 2 environmental MAGs from the Red Sea (MAG R02 GCA 023356605.1) and the San Francisco Bay (MAG GCA 040262915.1), (**Figure 1A**). Phylogenomic analyses of these new *Trichodesmium* sequences showed that Thieb has more diversity than previously defined and are divided into four groups/subclades (**Figure 1A**). Each new Thieb subclade was represented by at least 2 enrichment culture genomes and ≥1 genome per subclade also being sequenced from single filament isolated cultures (H9-4, VI-1, USCT24UA, USCT15UA, USCT02UA, and USCT12UA). These culture-defined subclades also clustered with MAGs sequenced from the Red Sea, San Francisco Bay, and the Atlantic Ocean, supporting representation of these strains both in culture and the global oceans (**Figure 1A**).

**Figure 1.**
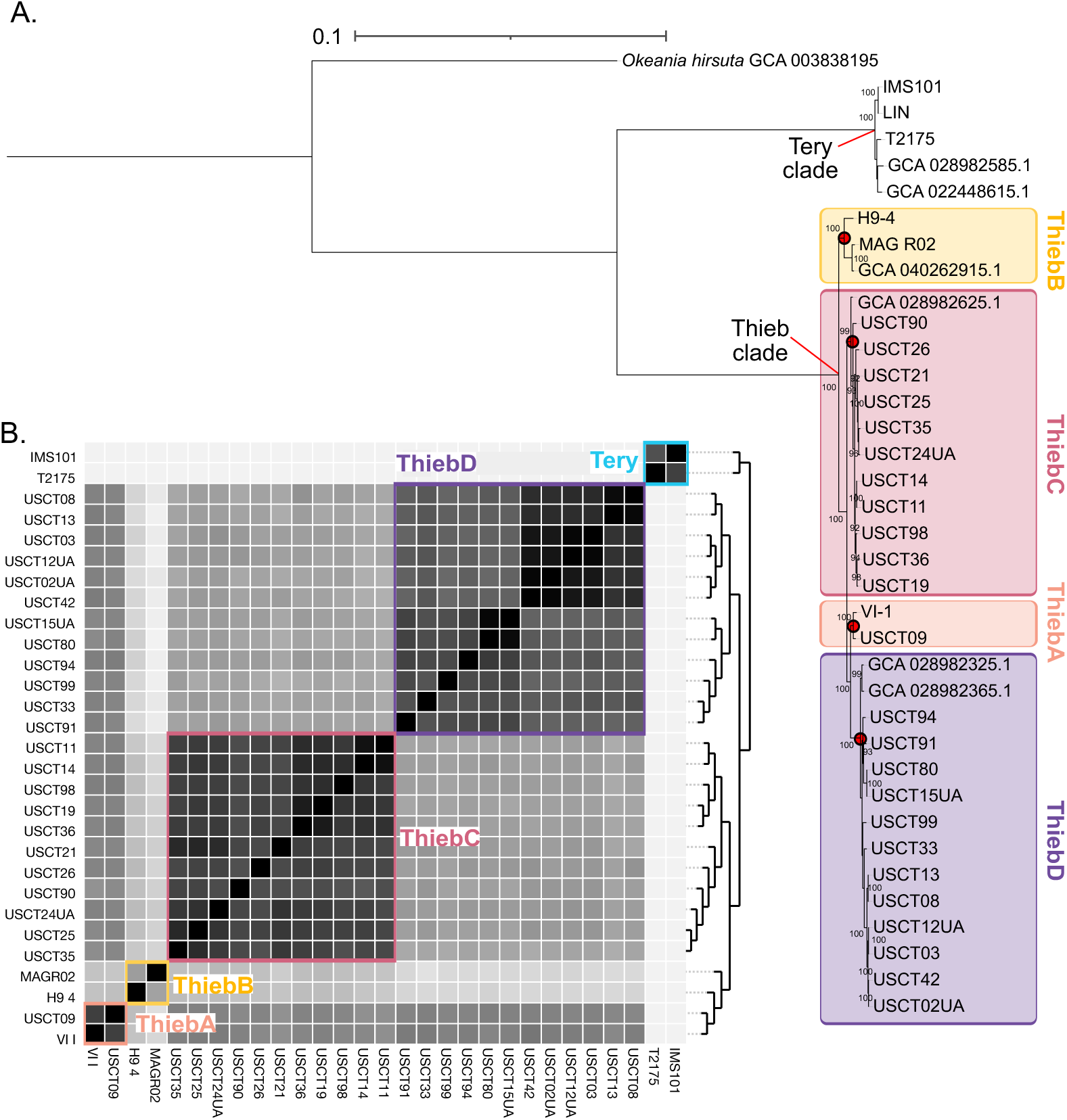
The phylogenomic tree of all *Trichodesmium* (A) rooted to *Okeania hirsuta*, and the average nucleotide identities (ANI) of all genomes (B) grouped by gene cluster frequencies (dendrogram shown to the right). The Thieb and Tery clades are denoted by a red line to each node, and the individual Thieb subclades are denoted by red dots at each node in the tree (A) and the color boxes in the tree and ANI heatmap (A, B). The lower threshold of the ANI in (B) is 97%.

Whole genome average nucleotide identities (ANI) of strain genomes were also used to further compare strain diversity (**Figure 1B**; **Supplemental Table S1**). At the ANI level, the same 4 groupings were present as in the phylogenomic tree (**Figure 1A-B**). However, the inter-subclade ANI values were found to all be >97%, which is above the predicted inter-species ANI threshold of 95% [18]. The difference between the Tery and Thieb clades, however, was ∼87%, which was more appropriate as a delineation between genomic species by ANI [18]. Some or all of these 4 Thieb subclades likely correspond to the botanical Thieb species, *T. thiebautii*, *T. hildebrandtii*, and *T. tenue*, which were historically identified by *hetR*, 16S-23S ITS phylogenetics and morphological differences [4, 14]. However, based on the highly conserved whole genome ANI values (∼97-98%), they were found to more accurately represent genomic subclades of clade Thieb rather than individual species. Therefore, these 4 new groupings within Thieb (**Figure 1A-1B**) were henceforth named ThiebA, ThiebB, ThiebC, and ThiebD rather than as separate botanical species (**Figure 1-B**).

### Subclades have taxonomically defined differences in colony morphology

Scanning electron microscopy (SEM) was used to explore more “macroscopic” colony morphological features and fine scale differences in cell dimensions. Representative strains from the most closely related subclades (**Figure 1A-B**), ThiebA and ThiebD, were used for SEM imaging of colonies (**Figure 2A-D**), dark field microscopy of colonies (**Figure 2E-G**), and individual filament imaging (**Supplemental Figure S1A-C**). These selected strains had also been initially purified from single filaments to ensure that enrichments were unialgal. ThiebD strain USCT33UA was found to form both tufts (**Figure 2A-B**) but also large, puff colonies with filaments aligned radially (**Figure 2E**). Contrastingly, the strain VI-1 in the closely related ThiebA subclade only formed smaller, hard to disassociate, puffs in culture (**Figure 2C-D**, **2F**) with filaments generally radiating out from the center. It was important to note that in **Figure 2C-D**, the ThiebA puff morphology was visualized in two puffs that had become attached, as noted by the pink arrow in the image. The un-attached “normal” morphologies for ThiebA colonies are shown in **Figure 2F**. Notably, ThiebA colonies had a dense matrix of material at the colony core. The boundaries of this matrix are noted with colorization in **Figure 2D**. Based on past imaging studies [59–61], this matrix of material was identified as *Trichodesmium* exopolysaccharides [62–64]. This dense matrix at the ThiebA puff colony core was a defined phenotypic difference from the more diffuse ThiebD tufts (**Figure 2A-B**) which appeared to have less EPS and a less dense colony core.

**Figure 2.**
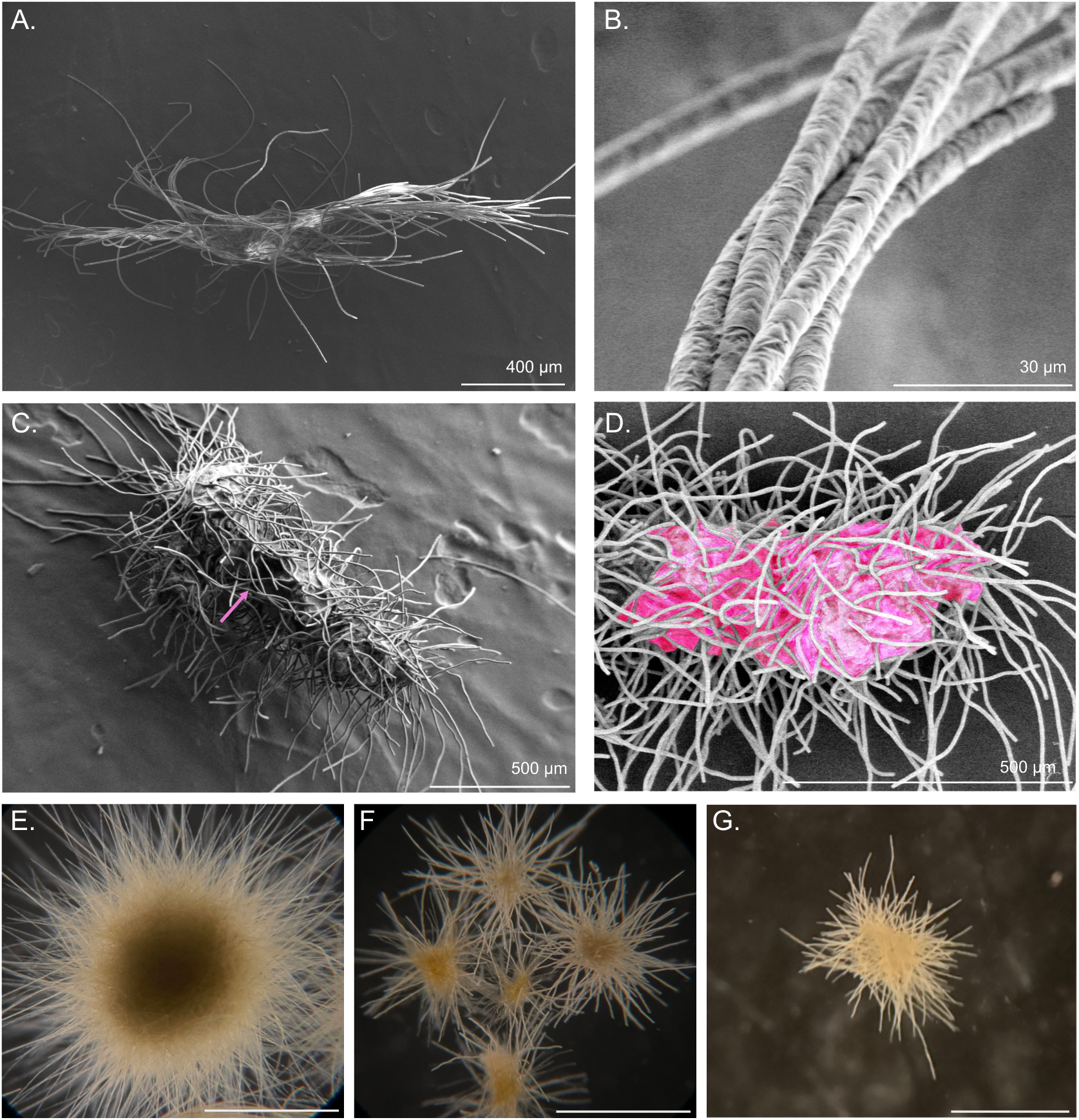
Morphological characterization of *Trichodesmium* subclades. Scanning electron microscopy (SEM) images taken of colonies of unialgal, representative strains of ThiebD USCT33 (A-B) and ThiebA VI-1 (C-D). The scale bars and sizes are listed on each image at the bottom. In (C), the pink arrow indicates the boundary between the two colonies that became attached. In (D) the exopolysaccharide “core” or the colony are colorized in pink to show the boundary. Dark field images of puff colony morphologies from ThiebD strain USCT33 (E), ThiebA strain VI-1 (F), and ThiebC strain USCT2P (G). The scale bars in E-G each indicate 1 mm.

Both puff and tuft shaped colonies are known to host a diverse suite of epibiont heterotrophic bacteria [65–70]. In the ocean, it has been demonstrated that associated epibionts and colony shape/size may likely influence micro-scale oxygen gradients. During nighttime, oxygen can be highest at colony peripheries and lowest at the center, where there is higher heterotrophic bacterial respiration and less diffusion with seawater [71–74]. In culture, unialgal ThiebA, ThiebC, and ThiebD strains were shown to develop very distinct colony morphology (**Figure 2**), despite being ∼97-98% ANI at the genomic level. Therefore, colonies belonging to these different Thieb subclades may also have differences in single colony oxygen gradients due to the denser EPS in the center of ThiebA puff colonies (**Figure 2C-D**), potentially allowing for higher heterotrophic bacterial respiration.

To closer examine these ecological dynamics caused by these different colony structures, higher magnification SEM imaging (**Figure 3**) was performed on the same colonies as in Figure 2A-D. When examining the EPS core at the center of the ThiebA puff colony (**Figure 3A-B**), a morphologically diverse and dense assemblage of associated epibiont bacteria were found physically embedded in the EPS matrix (which appeared as interwoven webs of polysaccharide strands and were initially identified from previous SEM images of EPS [75], (**Figure 3A-B**)). Phenotypically, these epibionts ranged from filamentous cells ∼2 µm in length by ∼100-200 nm in width, to rod shaped cells ∼1-2 µm in length and 200-400 nm in width (**Figure 3A-B**). Rod shaped bacteria ranged in shape from straight to curved (**Figure 3B**). GTDB-tk identification of MAGs assembled from this ThiebA VI-1 unialgal strain also indicated the presence of diverse heterotrophic epibiont genomes, and 4 of these MAGs even lacked a species designation from GTDB-tk (**Supplemental Table S3**). For comparison, high magnification images were also taken of ThiebD strain USCT33UA (**Figure 3C**), and these images show lower amounts of exopolysaccharides and bacterial cells. For example, a single, rod-shaped, bacterial cell (∼1 µm long by ∼200 nm wide) was found attached more on the periphery of the ThiebD filaments in the colony. EPS was also identified along ThiebD filaments (**Figure 3C**). However, the very high density of epibionts combined with a thicker EPS matrix at the core of ThiebA puff colonies (**Figure 2C-D**) supported the hypothesis that differences in epibiont density may explain the distinct oxygen gradients sometimes observed puff-shaped field colonies [74].

**Figure 3.**
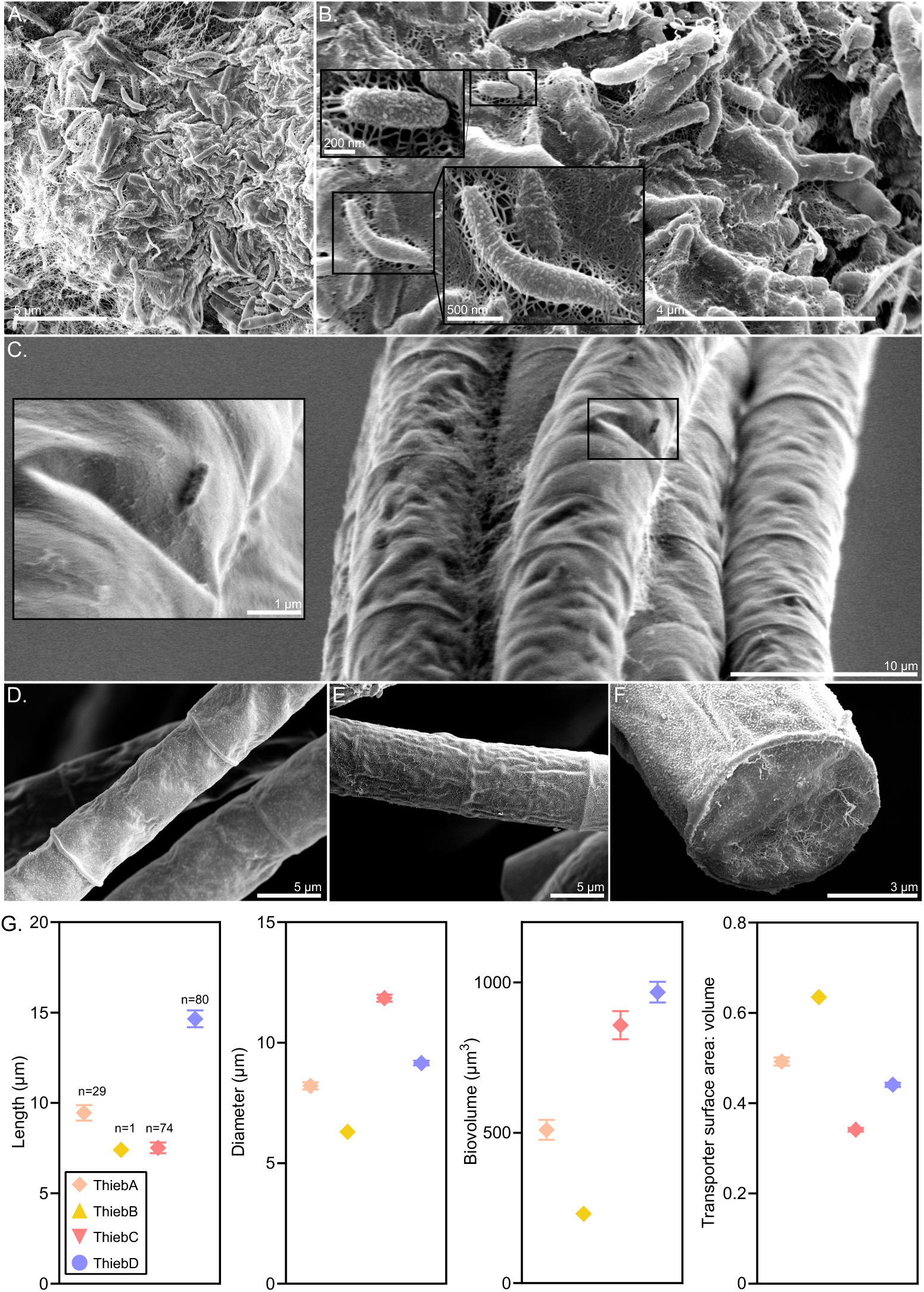
Scanning electron microscopy images taken at high magnification of epibiont heterotrophic bacterial cells adhered to *Trichodesmium* colonies. The ThiebA strain VI-1 exopolysaccharide colony “core” is shown in (A-B) with epibiont cells. ThiebD strain USCT33UA colony with an epibiont cell are shown in (C). Black boxes in (A-C) indicate sections where imaging was performed at lower (right boxes) and higher (left boxes) magnification to highlight specific cells/features. In (D), the ThiebD strain USCT33UA single cells within filaments are shown, and ThiebA cells are shown in (E-F). Panel (G) indicates measurements taken across strains and subclades with the y-axes indicating what measurement was taken for each plot. The subclades are indicated with different symbols and colors, and the “n=” values shown in the first plot indicate how many cells were measured across strains within each subclade.

### Intracellular biovolumes and surface area to volume ratios of Thieb are subclade-specific

High magnification SEM and phase contrast imaging also allowed for visualization of individual *Trichodesmium* cells in addition to epibionts. ThiebD cells in filaments (**Figure 3D; Supplemental Figure S1A**) were longer than both ThiebA (**Figure 3E; Supplemental Figure S1B)** and ThiebC (**Supplemental Figure S1C**). Two notable, distinguishing characteristics of ThiebA and ThiebC were that the former appeared to have “covered” or less visible septa (**Figure 3E**, **Supplemental Figure S1B**), and cells in the latter were consistently much smaller in length than in width (**Supplemental Figure S1C**). In initial characterizations of the botanical species of *Trichodesmium* in the late 1800’s, *T. thiebautii* were noted to have covered septa, and *T. hildebrandtii* were noted to have a diameter 3x as large as the length [14]. Therefore, while these new strains cannot be definitively matched to botanical classifications at the gene-centric level, ThiebA and ThiebC shared the most morphological similarity with *T. thiebautii* and *T. hildebrandtii*, respectively. Another notable feature was that ThiebA filaments appeared to have defined “end caps” in this differentiated cell type (**Figure 3F**). Consistent with whole colony images, individual EPS strands were also adhered to this ThiebA terminal cell (**Figure 3F**).

In addition to SEM, phase contrast imaging was performed on live cells that had not been fixed or dehydrated. Cell lengths and diameters were measured for ≥2 strains in each subclade, with the exception of ThiebB for which there were cell measurements taken for only one strain [4, 39], and measurements were used to calculate biovolumes and transporter surface area to volume ratios (SA:V). ThiebD had the longest cells while the shortest cells were observed in ThiebB and ThiebC (**Figure 3G**). However, ThiebC had the largest diameter. As a result, both ThiebD and ThiebC had large biovolumes, but it was a result of different cellular dimensions (**Figure 3G**). ThiebB and ThiebA, being the smallest overall, had the largest transporter available SA:V, whereas ThiebD and ThiebC had the lowest (**Figure 3G**). Therefore, despite these subclades being isolated from generally low nutrient locations, these findings suggest that ThiebA and ThiebB are likely more optimized for somewhat lower nutrient regimes, while ThiebD and ThiebC are likely more and optimized for somewhat higher nutrients [76]. Also, depending on which of these subclades dominates the oceanic *Trichodesmium* community at any given time, there may be differences in *Trichodesmium* C+N standing stocks because of these varied subclade-specific biovolumes.

### Genomic characteristics and subclade-specific accessory genes reveal distinct metabolic niches

Detailed Thieb clade pangenome analyses were performed in Anvi’o [33, 50] to determine if subclade designations at the cell morphology, whole genome and ANI level corresponded to specific differences in gene content. The 25 new genomes from this study alongside 3 previously sequenced culture genomes (strains IMS101 and 21-75 in the Tery clade and Thieb strain H9-4 [17, 77], and one environmental ThiebB MAG from the Red Sea [10] were included in the pangenome. The pangenome was ordered by gene cluster presence/absence across all strains, and an MCL of 10 was used to determine the threshold of homologous gene clusters (See: **Materials and Methods**). Visualization of the sorted pangenome revealed that the largest grouping of shared, single copy gene clusters was in the genomic core bin, which included 1,862 gene clusters found in all strains (**Figure 4A**). These included the genes involved in photosynthesis, nitrogen fixation, and the other basal metabolic characteristics of *Trichodesmium* (**Supplemental Table S6**).

**Figure 4.**
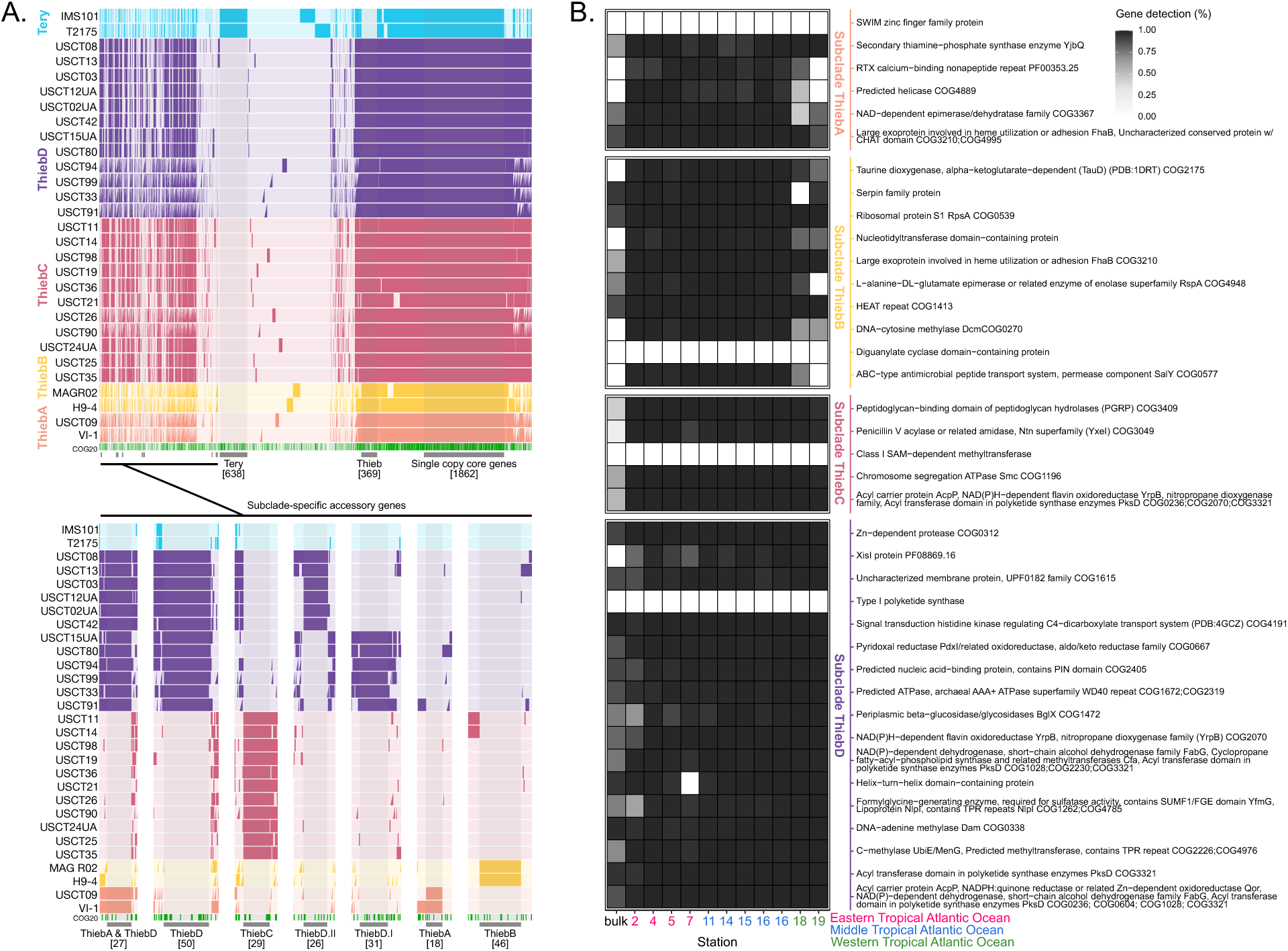
The presence/absence pangenome of Thieb alongside 2 Tery strains (A) and the gene-centric detection of subclade specific accessory genes (B). In (A), each strain is shown across rows, ordered by gene cluster frequency, and colored according to the labeled subclade (left). The top panel indicates the entire pangenomic alignment, and the bottom panel indicates a zoomed in plot of just the subclade specific accessory genes aligned to the left in the full pangenome. The gray bins and shading on the pangenomic alignment are labeled according to which clade/subclade the gene clusters belong to and how many gene clusters there are in each bin (i.e., the “Tery” bin indicates the 638 gene clusters only found in Tery, the “Thieb” bin are 369 found only in Thieb, etc.). The annotated accessory genes in the bottom panel of (A) are shown again in (B) with their environmental % detection in ocean metagenomes collected across and east to west transect in the Atlantic Ocean. The % detection (0-100%) in (B) indicates how much of the individual genes from cultures of each subclade were also sequenced in metagenomic samples (i.e., a value of 100% means the whole gene sequence from a given culture was found in the metagenome). The stations for the transect are given at the bottom with the color indicating geographic location (B). The station labeled as “bulk” originated from whole community plankton rather than just from *Trichodesmium* colonies like the other samples.

Within the major *Trichodesmium* clades, there were ∼1.7x as many unique gene clusters (638 clusters) shared by all Tery vs those shared by all Thieb (369 clusters), (**Figure 4A**). Only ∼44% of the Tery-specific genes in strain IMS101 had a known function with NCBI-COGs, but of the genes that were annotated, some notable ones included accessory copies of the sulfate transporter CysZ (PDB:3TX3) annotated by NCBI-COGs, Pfam, and KOfam and an additional copy of *feoA* (PDB:2GCX) predicted to be involved in Fe^2+^ transport (**Supplemental Table S6**). Many of the other Tery accessory genes were overall related to NCBI-COG-annotated mobilome, signal transduction mechanisms, cell wall/membrane/envelope biogenesis, and secondary metabolite biosynthesis. Comparatively, in the Thieb accessory genes in strain H9-4, ∼55% of the gene clusters were annotated by NCBI-COGs. Within these gene clusters, many genes were transposase/mobilome related. There were also additional accessory copies of genes such as *proA* gamma-glutamyl phosphate reductase (PDB:4GHK) likely involved in generation of proline from glumate [78], alkaline phosphatase D (*phoD*), (PDB:2YEQ), two copies of Fe^2+^ transport gene *feoA* (PDB:2GCX), and one copy of Fe^2+^ transport *feoB* (PDB:3TAH), (**Figure 4A**). The presence of accessory *feo* genes in both Thieb and Tery suggested that these proteins diverged enough in sequence such that they did not align, even with a pangenome MCL inflation value of 10 used for highly similar genomes [33, 40, 50]. As well, additional, accessory copies of the predicted alkaline phosphatase *phoD* in Thieb may be indicative of biogeographic differences between Thieb vs Tery wherein Thieb are generally more abundant in offshore, phosphorus-limited, oligotrophic oceans, like summertime in the Sargasso Sea [8, 9, 79, 80].

Within the Thieb clade, the subclades ThiebA, ThiebB, ThiebC, and ThiebD each contained ecologically relevant subclade-specific accessory gene clusters. As shown in **Figure 4A**, ThiebD had the most accessory gene clusters (50 clusters) followed by ThiebB (46 clusters), ThiebC (29 clusters), and ThiebA (18 clusters). There were also accessory clusters found in other groups, such as the 27 shared clusters between ThiebA and ThiebD (**Figure 4A**), which were the two most closely related subclades (**Figure 1A**). These 27 shared genes included genes related to fatty acid biosynthesis, toxin-antitoxins, and post translational modification (**Figure 4A**; **Supplemental Table S6**). Other notable bins of shared gene clusters included ThiebD strains in bin “ThiebD.I,” which were isolated from the Bermuda Atlantic Time Series (BATS) site over multiple years and had 31 accessory clusters, particularly the *wcaK* gene (PF04230.19) involved in exopolysaccharide biosynthesis [81, 82] and the glycosyltransferase *rfaJ* (PDB:1G9R) involved in lipopolysaccharide biosynthesis (**Figure 4A**), [83, 84]. Also, within ThiebD, another bin called “ThiebD.II” contained 26 gene clusters from strains only isolated from the gulf stream (general location: offshore east of Ocean City, Md), (**Figure 4A**; **Supplemental Table S1**). Strains contributing to these ThiebD.II gene clusters were enriched from a single sampling day/time and therefore may have come from a near clonal bloom of *Trichodesmium*, although they were sequenced from both puff and tuft colonies (**Supplemental Table S1**). Contrastingly, strains contributing to the ThiebD.I bin were collected across multiple seasons and even years from BATS. Therefore, even within the subclade (ThiebD), there were biogeographically constrained differences in accessory gene clusters which potentially contribute to ecological function. Specifically, differences in exopolysaccharide biosynthesis genes between Thieb strains may affect overall contributions to carbon cycling. Further sequencing ThiebD isolates from the gulf stream and other regions and comparison to ThiebD MAGs assembled from environmental datasets will help to resolve these genomic trends and inform on ecological function of ThiebD across the oceans.

While these accessory genes are present in genomes sequenced from cultures, presence or absence of genes does not immediately inform on importance in the environment. Therefore, accessory genes from each Thieb subclade were used as read recruitment targets to a *Trichodesmium* metagenomic dataset wherein *Trichodesmium* colonies were collected in plankton nets, separated from the bulk plankton community with serological pipettes, sorted by puff or tuft morphology, filtered down, and later sequenced [17]. This was done across a transect from the eastern to the western tropical Atlantic Ocean. Metagenomic short reads from this dataset were then mapped to Thieb accessory genes from representative unialgal strains from each subclade (**Figure 4B**). Gene-centric detection was used to determine whether individual genes sequenced from cultures had also been sequenced in the environment. From this, it was found that most of the subclade-specific genes were all detected across all or most stations, apart from the control (Bulk column; **Figure 4B**), which was a bulk net tow of the entire plankton community and contained relatively less *Trichodesmium*. Genes that weren’t detected at all (or potentially had DNA sequences too short to be detected) included a SWIM zinc finger, diguanylate cyclase, class I SAM−dependent methyltransferase, and type I polyketide synthase gene. However, most accessory genes were detected, and there were biogeographical trends by subclade. On the eastern equatorial Atlantic side of the transect, the detection values for the ThiebD-only accessory genes were generally lower whereas the other 3 subclades were generally higher. However, on the western equatorial Atlantic side, detection values of ThiebD accessory genes were generally higher whereas ThiebA and ThiebB genes dropped to lower or even 0% detection (**Figure 4B**). This revealed that subclades likely had unique biogeographic signatures during this sampling, and while all subclades were generally present, there were likely differences in abundances of each.

It is noteworthy that several accessory genes across subclades were specialized metabolite related (e.g., such as the acyl carriers, acyl transferases involved in polyketide synthase, and taurine dioxygenase). Specialized metabolites likely structure epibiont communities in cyanobacteria, including *Trichodesmium* [85, 86]. Thus, the overall specialized metabolite-related genes in all Thiebs were further examined using AntiSMASH 8.0 [44] for annotation of biosynthetic gene clusters (BCGs), and BiGSCAPE 2.0 [45] was used for clustering and taxonomic placement. Overall, Thieb genomes were most enriched in BCGs related to non-ribosomal peptide synthetases (NRPSs), terpenes, and type I polyketide synthases, and representative genomes from each subclade had 18-23 BCGs involved in some form of specialized metabolite synthesis (**Supplemental Table S7; Supplemental Figure S2A).** There was one biosynthetic gene cluster that fell into two BigSCAPE similarity family clusters, was present in all Thieb genomes, and had the closest AntiSMASH similarity match to a Type I NRPS responsible for synthesizing hexose-palythine-serine/hexose-shinorine (**Supplemental Figure S2B**). These specialized metabolites have been identified as mycosporine amino acids [87], which can be produced by cyanobacteria and help to prevent cellular damage from incoming UV light in the ocean euphotic zone [88]. Therefore, this gene cluster in Thieb is likely an adaptation for highly stratified, oligotrophic regions like the Sargasso Sea in summer seasons wherein the incident light from the sun is most intense [89, 90]. Also present in nearly all Thiebs (24/27 genomes) and across all subclades was a biosynthetic gene cluster in two BigSCAPE similarity family clusters which had high AntiSMASH similarity confidence to a cluster responsible for synthesis of the alkene, 1-heptadecene (**Supplemental Figure S2C**). This compound (C_17_H_34_) is a hydrocarbon which cyanobacteria produce from fatty acids [91], and the Tery strain IMS101 has been demonstrated to produce the alkane, heptadecane [91]. For Thiebs, production of long hydrocarbons would likely contribute to carbon cycling inside single colonies and provide heterotrophic epibionts with a rich carbon source in an otherwise low carbon ocean region.

The biosynthetic gene clusters in all or subclusters of Thiebs likely contribute to the ecological structure of the broader clade in the oligotrophic oceans. However, as discussed previously, there were also specialized metabolite-related genes present only in specific subclades. In the AntiSMASH data, many subclade-constrained biosynthetic gene clusters lacked a known function or had low similarity confidence to known clusters. Some of these included region 470.1 in USCT24UA (**Supplemental Table S7**) that was present 9/11 of ThiebC strains, region 565.1 in USCT12UA (**Supplemental Table S7**) present in 6/12 ThiebD genomes and the 2 ThiebA genomes, and region 475.1 in USCT24UA (**Supplemental Table S7**) present in 7/11 ThiebC genomes with low similarity confidence to a gene cluster for desmamide A/desmamide B/desmamide C synthesis. This product is a lipoglycopeptide that has been demonstrated as a probable anti-microbial molecule [92]. There was also one biosynthetic gene cluster identified only in ThiebD and a partial cluster in ThiebB strain H9-4 that had medium AntiSMASH similarity confidence in ThiebD strains to a BGC involved in heterocyst glycolipid formation (**Supplemental Figure S2D**), despite *Trichodesmium* not forming heterocysts. Recently, it has been demonstrated that non-heterocystous cyanobacteria can still have heterocyst glycolipid genes that are likely involved in diverse processes besides O_2_ protection in heterocysts [93].

Differences in specialized metabolite production across subclades are ecologically relevant as they are likely to change which organisms are utilizing and cycling *Trichodesmium*-derived carbon and nitrogen. *Trichodesmium*-derived C+N helps form the base of oligotrophic gyre food webs through their fixation of both CO_2_ and N_2_ gas [1], so understanding what specific compounds they synthesize and how these compounds may affect associated organisms or grazers is essential to understanding C+N cycling in oligotrophic ocean regions.

### Species-level relative abundances differ in whole water and *Trichodesmium* colony metagenomic samples

Individual gene-centric detection of accessory genes from each Thieb subclade differed between *Trichodesmium* samples collected from the western vs eastern equatorial Atlantic Ocean (**Figure 4B**). Therefore, to explore how subclade abundances changed across this same transect, total mean coverage values of subclade-specific accessory genes were also compared. Overall, on the eastern side of the equatorial Atlantic, there were only puff colonies collected, and samples were dominated by ThiebA and ThiebC (**Figure 5A**). This was in line with lab-based culturing wherein ThiebA and ThiebC both formed puffs (**Figure 2F-G**) and were isolated from almost entirely puff shaped colonies (**Supplemental Table S1**). In the middle equatorial Atlantic, samples were generally dominated by a mixture of all 4 subclades and there was the highest relative abundance of ThiebB, which can form puffs or tufts (**Figure 5A**; **Supplemental Table S1**). To the western equatorial Atlantic, ThiebD were most abundant in the metagenomic reads collected from a bulk plankton net tow (although the overall *Trichodesmium* abundance was lower in this sample) and in the tuft colonies that were collected (**Figure 5A**). This region was also where the ThiebD MAGs, GCA 028982325.1 and GCA 028982365.1 (**Figure 1A**), were assembled from [17]. This trend was also consistent with cultures of ThiebD that formed tufts (**Figure 2A-B**) but also formed loosely associated puffs (**Figure 2E**) and were enriched from both puff and tuft colonies (**Supplemental Table S1**). Therefore, overall, environmental abundances of individual subclades can be directly linked to colony morphologies in culture. Importantly, colony morphology and shape have also been proposed as a determining factor in epibiont bacterial community structure within *Trichodesmium* colonies [69], which can therefore affect oxygen concentrations and *Trichodesmium*-derived C+N biogeochemical cycling.

**Figure 5.**
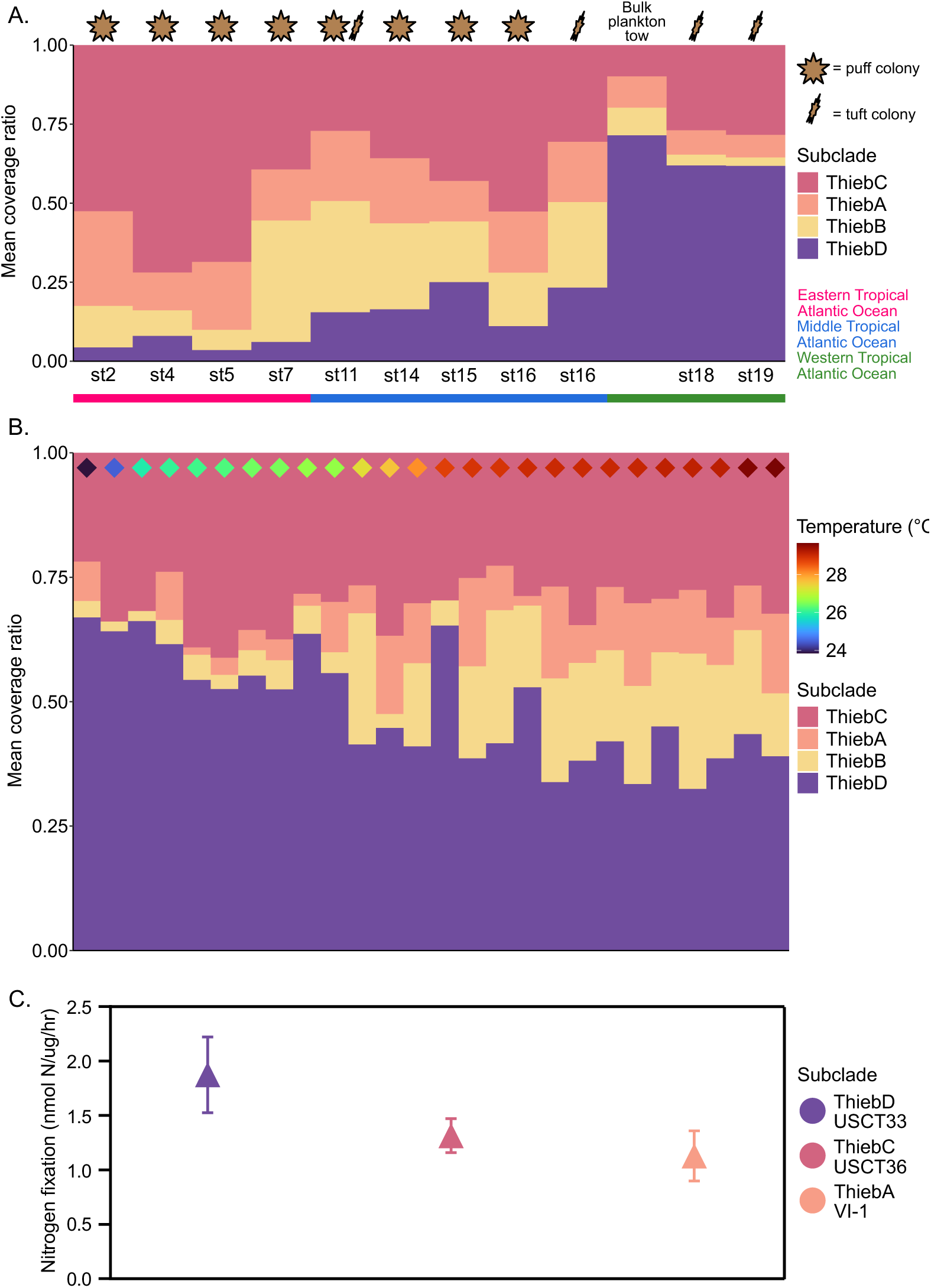
Read recruitment mean coverage ratio (relative abundance) results for two oceanic metagenomic datasets and nitrogen fixation and growth rates of Thieb subclades. Mean coverage indicates the average read coverage across the accessory genes for a representative strain from each subclade. In (A), the abundances of subclades in *Trichodesmium* picked-colony metagenomes collected across the eastern to western equatorial Atlantic are shown. The type of colonies that were used to generate the metagenomic samples from each station is indicated by the “puff” or “tuft” symbol at the top. In (B), the relative abundances of Thieb subclades are shown across bulk water (>0.2 µm-filtered) metagenomic samples and the x-axis was sorted by temperature. The station temperature where each sample was collected is also indicated by the color of the diamond symbols at the top of each plot. Chlorophyll-normalized nitrogen fixation values for select enrichments are shown in the first panel of (C).

In addition to picked-colony metagenomic reads, the GO-SHIP metagenomic reads [47] collected from 0.2 µm-filtered whole seawater samples, were used to explore bulk water *Trichodesmium* community dynamics. Metagenomic reads were read-mapped to subclade-specific accessory genes and relative mean coverage (abundance) values were compared to the temperatures of each station. In stations where at least one Thieb subclade was at >1x mean coverage, ThiebD was at highest relative abundance in the cold-water stations (**Figure 5B**). However, at temperatures of >28 °C, there was a relative abundance shift. In these samples, ThiebD still composed ∼30-50% of the total Thieb community, but ThiebA and ThiebB became relatively more abundant. To explore this deeper, Thieb abundance patterns at each station were compared in an NMDS using Bray-Curtis distances, UMAP, and k-means clustering (**Supplemental Figure S3A-B**). The community composition at each station plotted in the NMDS (Stress = 0.04) corresponded to two overall k-means clusters, and these clusters generally corresponded to high (>28 °C) vs lower (<28 °C) station temperature (**Supplemental Figure S3A**). The exception to this was 4 high temperature stations (out of 26 total stations) that instead clustered with the colder samples. This could potentially be because ThiebD was generally the most abundant subclade at all stations in GO-SHIP but decreased relatively with increasing temperature. Additionally, it could be due to other variables, such as iron or phosphorus, that also affected biogeographical patterns in this transect. The warmest stations where Thieb was present in the GO-SHIP transect were proximal to the Amazon River plume, which potentially also influenced the distribution of *Trichodesmium* [94]. However, the temperature gradient and biogeographical trends generally aligned with culturing thermal optima; when grown at a “colder” (<28 °C) temperature of 26 °C, a representative strain of ThiebD had a faster nitrogen fixation rate than ThiebA strain VI-1 and ThiebC strain USCT36 (**Figure 5C**). As well, these trends do agree with predictions that warming oceans will favor smaller phytoplankton [95], such as ThiebA and ThiebB.

These environmental abundance values combined with lab cultivation and physiology revealed that, as temperatures continue to warm under climate change conditions, we may see a Thieb community shift towards more ThiebB (and somewhat ThiebA), particularly relative to ThiebD. This shift could potentially lead to lower C+N standing stocks of Thieb in the global ocean as ThiebB also had the smallest intracellular biovolume and ThiebD had the highest. As well, due to Thieb subclades having different SA:Volume ratios, shifting concentrations of phosphorus and iron in the oligotrophic oceans in a changing global climate may cause additional biogeographical differences. As Thieb can be found to be ∼90% of the whole *Trichodesmium* community [8, 9], an overall shift in Thieb subclade dynamics could have significant implications for *Trichodesmium*’s contribution to C+N biogeochemical cycling and export in the oligotrophic oceans. Future isolation efforts focused on ThiebB and simulated climate change culturing experiments will help to resolve these trends and allow for predictions of how clade Thieb will respond to increasing ocean temperatures.

## Conclusion

*Trichodesmium* provide a localized source of bioavailable C+N in oligotrophic gyres [1], which make up a majority of the global surface oceans [20]. Through the conversion of N_2_ and CO_2_ gas into organics and NH_3_, *Trichodesmium* colonies can each be thought of as a micro-scale “oasis” of resources that other organisms such as zooplankton grazers and epibiont bacteria can utilize [66, 96]. However, as demonstrated herein, Thieb colonies, which comprise ∼90% of the total *Trichodesmium* cells in the ocean [8, 9], can be very different depending on which subclade they correspond to. Despite being ∼97% ANI, Thieb subclades were demonstrated to form different colony shapes, with ThiebD and ThiebB growing as both tufts, puffs, and free filaments vs ThiebA and ThiebC, which generally just form tightly packed puffs. Also, ThiebA puffs appeared to form a dense EPS core at the center of their colonies, whereas ThiebD colonies appeared looser with less EPS. These differences would likely structure *Trichodesmium*-associated epibiont communities differently, may lead to increased respiration rates within EPS-rich colonies, and overall, potentially cause differences in micro-scale O_2_ gradients [74].

In addition to colony morphology, Thieb subclades also had differences at the cellular level. ThiebD and ThiebC had the largest biovolumes and lowest SA:volume, and ThiebB and ThiebA had the smallest biovolumes and highest SA:volume. This was ecologically relevant as ThiebD had the highest relative abundance at colder temperature stations in 0.2 µm-filtered GO-SHIP [47] seawater metagenomes and lower relative abundance at high temperature. At stations where sea surface temperatures were >28°C, ThiebB (and slightly ThiebA) became relatively more abundant. Under climate change warming where sea surface temperatures may frequently reach >28°C in gyres, phytoplankton abundances may shift towards smaller cells [95, 97]. Therefore, there may be corresponding shifts in Thieb-associated C+N standing stocks as the *Trichodesmium* communities potentially shift towards smaller-celled ThiebB and possibly ThiebA. This has implications for bottom-up control in oligotrophic oceans where *Trichodesmium* often contributes to the structure and extent of N (and sometime C) cycling and export from the surface oceans.

In addition to taxonomically constrained differences in biovolume-predicted estimates of total C+N content in Thieb subclades, different Thieb subclades had distinctly conserved accessory gene content that could influence epibionts and community structure. For example, Thieb subclades each had specific unique genes, some of which were involved in specialized metabolite biosynthesis and a large majority which were detectable in environmental samples as well as in cultures. Changes in specialized metabolite production would also likely structure epibiont communities differently and therefore influence *Trichodesmium*-derived C+N biogeochemical cycling in the global oceans.

Prior to this study, there was only one whole genome sequence from a culture of Thieb, and only 1 extant Thieb culture. Herein, 25 new culture genomes were sequenced alongside their epibionts, and these genomes were used to link culture morphology, physiology, biogeography and functional capability in the environment. It was found that, while genetically similar, Thieb subclades are likely to have meaningful differences in lifestyle that structure the overall contribution of *Trichodesmium* to marine C+N cycling. Also, if the climate continues to shift, we may see differences in subclade abundances caused by rising temperatures, increasing stratification in ocean gyres, and shifts in sea surface nutrients. By leveraging these cultures and genomes, future studies can now further explore how the physiology of these organisms will change in response to climate change-simulated differences in nutrients (iron and phosphorus), temperature, and different light fields.

## Acknowledgements

We thank the crew of the R/V Atlantic Explorer and its crew as well as the scientific team of the Bermuda Atlantic Time Series. We thank Rod Johnson and Claire Medley for their support with sample collection and field work. We also thank undergraduate students Adam Wadhwani, Max Binkowski, and Nicolette Lee with assistance with lab-based culturing of *Trichodesmium*.

Funding was supported by the US Nation Science foundation grant 2125191 to EAW.

## Contributions

EAW – field work sampling, culture isolations

CSC – single colony and filament isolations, SEM and phase contrast imaging, genomic analyses

AMB – field work sampling and sample collection, culture collection

YZ – sequenced VI-1 genome

SJB – filament isolations of lab strains, field work sampling

All - contributed to writing and/or editing

---

**Figure S1.**
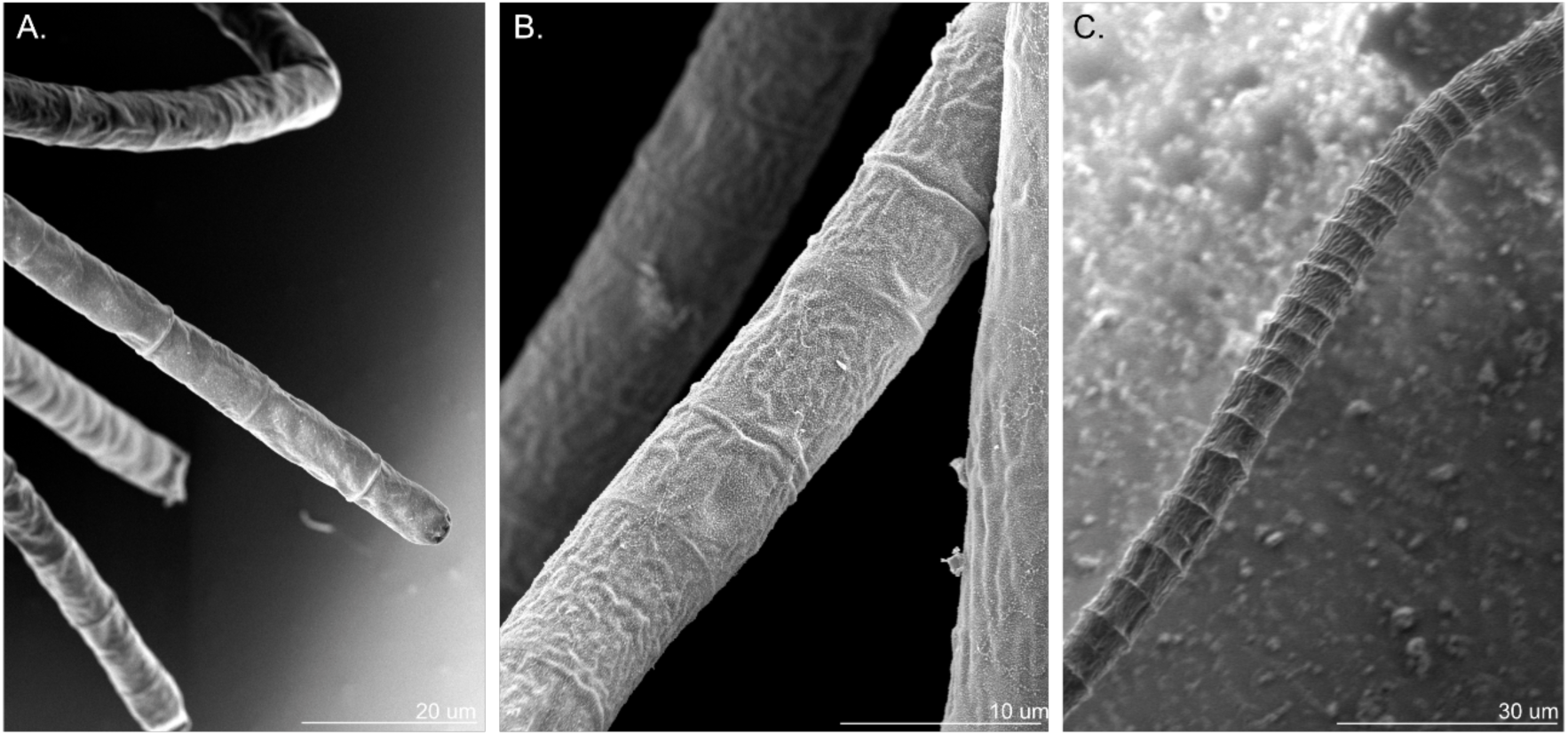
Scanning electron images show single filaments of ThiebD USCT33 (A), ThiebA VI-1 (E), and ThiebC USCT36 (F). The scale bars with sizes are indicated on each image.

**Figure S2.**
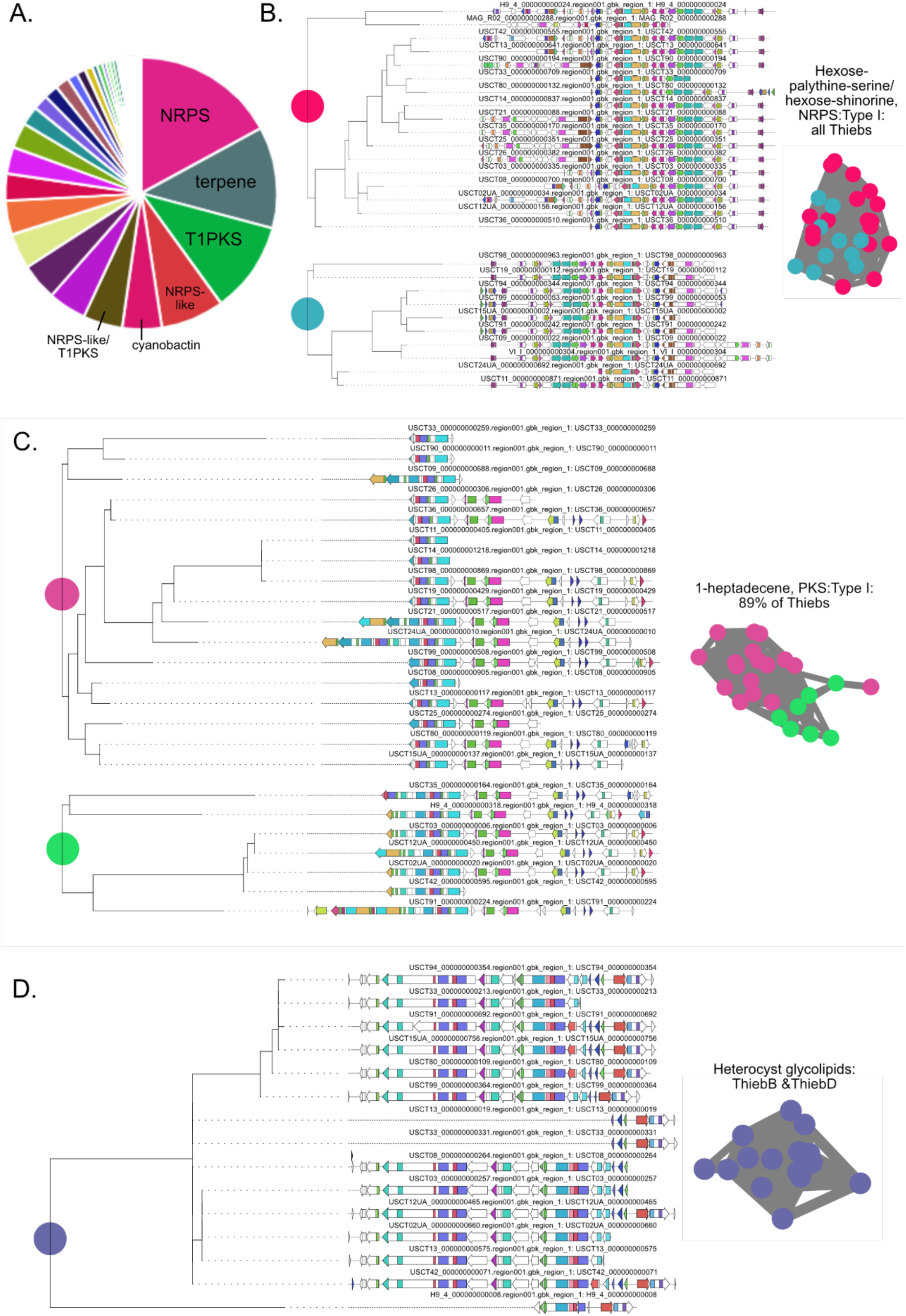
Biosynthetic gene clusters (BCGs) and taxonomic analysis of specialized metabolite genes across Thieb. In (A), the pie chart shows the types of different specialized metabolite categories for BCGs that are enriched across all Thieb genomes in the analysis. In (B-D), the alignment and phylogenetic grouping of BCGs for different metabolites (named to the right of each tree) are shown. The BiGSCAPE clusters that the BCGs fall into are indicated by the colored dots at the nodes of each tree, and the clusters are also plotted to the right of the tree with each dot in the cluster indicating 1 strain and the colors of the dots denoting each cluster. In (B-C), strains made the same compound but still had 2 distinct overlapping BiGSCAPE clusters for each compound as the BGCs were divergent in gene sequence across strains.

**Figure S3.**
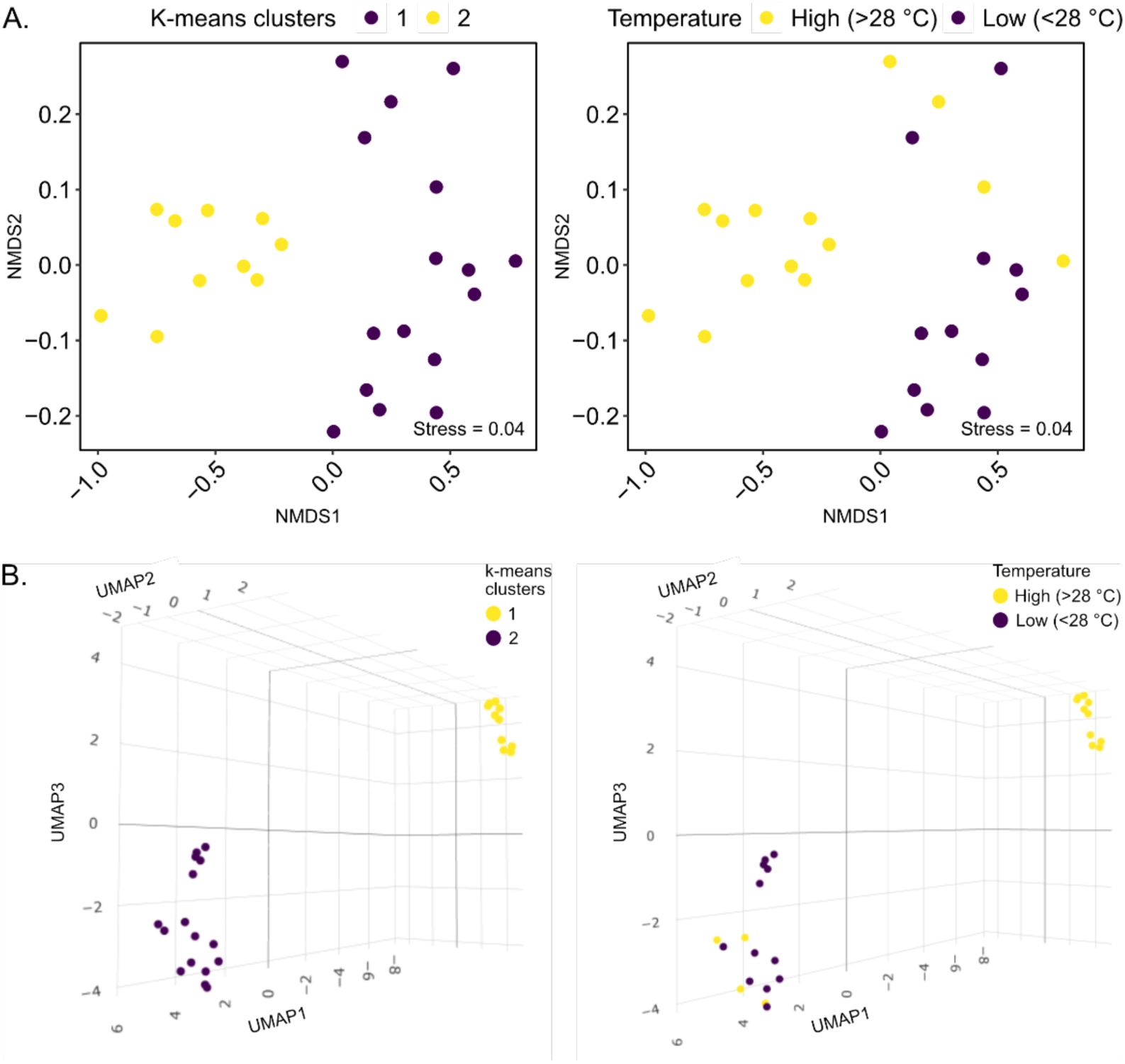
The NMDS/Bray-Curtis values of Thieb abundances with k-means clustering across the GO-SHIP bulk water samples where *Trichodesmium* were found (left panel) as compared to station temperature on the same plot in the right panel (A). In (B), the UMAP and k-means clustering of Z-scored Thieb abundance values from the same dataset are shown in the left panel as compared to station temperature in the right panel.

